# Microbial community composition and diversity via 16S rRNA gene amplicons: evaluating the Illumina platform

**DOI:** 10.1101/010058

**Authors:** Lucas Sinclair, Omneya Ahmed Osman, Stefan Bertilsson, Alexander Eiler

**Affiliations:** Department of Ecology and Genetics, Limnology, and Science for Life Laboratory, Uppsala University, Uppsala, Sweden

## Abstract

As new sequencing technologies become cheaper and older ones disappear, laboratories switch vendors and platforms. Validating the new setups is a crucial part of conducting rigorous scientific research. Here we report on the reliability and biases of performing bacterial 16S rRNA gene amplicon paired-end sequencing on the MiSeq Illumina platform. We designed a protocol using 50 barcode pairs to run samples in parallel and coded a pipeline to process the data. Sequencing the same sediment sample in 248 replicates as well as 70 samples from alkaline soda lakes, we evaluated the performance of the method with regards to estimates of alpha and beta diversity.

Using different purification and DNA quantification procedures we always found up to 5-fold differences in the yield of sequences between individually barcodes samples. Using either a one-step or a two-step PCR preparation resulted in significantly different estimates in both alpha and beta diversity. Comparing with a previous method based on 454 pyrosequencing, we found that our Illumina protocol performed in a similar manner – with the exception for evenness estimates where correspondence between the methods was low.

We further quantified the data loss at every processing step eventually accumulating to 50% of the raw reads. When evaluating different OTU clustering methods, we observed a stark contrast between the results of QIIME with default settings and the more recent UPARSE algorithm when it comes to the number of OTUs generated. Still, overall trends in alpha and beta diversity corresponded highly using both clustering methods.

Our procedure performed well considering the precisions of alpha and beta diversity estimates, with insignificant effects of individual barcodes. Comparative analyses suggest that 454 and Illumina sequence data can be combined if the same PCR protocol and bioinformatic workflows are used for describing patterns in richness, beta-diversity and taxonomic composition.

## Introduction

As the dropping price of DNA sequencing pushes laboratories and core facilities to switch from one established method to another, it is critical to check the validity and effects of the newly adopted processes. Researchers are often forced by this fast-paced evolution of technology to use the latest machines while abandoning the old ones, all too often neglecting that the correct setup and validation of such methodology is a crucial step in conducting rigorous scientific research. We would like to share our experience and the lessons learned from using the Illumina MiSeq platform as our solution for performing amplicon sequencing, replacing the established 454 pyrosequencing protocol previously used in our laboratory.

Amplicon sequencing, in particular that of the small subunit rRNA gene (16S rRNA gene in Bacteria and Archaea or 18S rRNA gene in Eukarya), is a widely applied approach to study the composition, organization and spatiotemporal patterns of microbial communities, due to its ubiquity across all domains of life (1). In the last decades, 16S rRNA gene amplicons were analyzed using fingerprinting techniques such as TRFLP (2) and ARISA (3) in combination with clone library construction and Sanger sequencing. However, this often provided insufficient coverage to describe and compare microbial communities (4). Now, high-throughput sequencing (HTS) technology and the application of barcode indexing are allowing the collection of thousands of sequences from a large number of samples simultaneously (5) (6). These approaches have revealed deeper insights into the diversity of microbial communities (7) (8) and, by increasing sample numbers, have expanded the possibilities to study community and population dynamics over much finer temporal (9) and spatial scales (8).

Still, the presence of sequencing errors and base miscalling together with PCR errors, chimera formation and pseudogenes introduce noise thus biasing estimates of diversity and taxon abundances. These concerns have been elaborated in great detail previously (10) (11) (12) (13) (14) (15) (16), and together, these studies suggest that HTS based surveys require substantial data denoising.

Amongst these HTS technologies, Illumina is currently the state of the art when it comes to 16S rRNA gene amplicons (17) (18) (19) (20) (21) (22) (23). It surpasses the previous 454 technology most notably by its price difference, coming in at about 0.5 USD/Mb for a MiSeq installation against 31 USD/Mb for a 454 GS Junior installation (24).

Here, we evaluated the methods suggested in these publications and provide our view on best practices for following the dynamics of microbial taxa as well as for estimating alpha and beta diversity. Alpha diversity estimates tested include Chao1 and ACE for richness, Pilou’s, Shannon Wiener’s and Simpson’s evenness estimates. Beta diversity estimates were based on weighted UniFrac distances and Bray-Curtis dissimilarities. We described a procedure starting with PCR amplification of bacterial 16S rRNA genes, paired-end Illumina sequencing and ending with bioinformatic analyses.

## Materials and Methods

### Collection

A sediment sample was collected from the highly phosphorus-saturated sediments of Lake Vallentunasjön (N59°29’24” E18°01’42”) in Sweden between 1-2 cm of depth. The sediment core was incubated at 21 °C prior to sampling (25). In addition, 9 samples from 3 Austrian soda lakes located on the eastern bank of the Neusiedlersee (N47°44’ W16°49’) were obtained from the water column during the winter of 2012 through summer of 2013 by capturing cells on 0.2 μm filters. Samples were kept frozen at -80 °C. Nucleic acids were extracted using the Powersoil DNA isolation Kit (MO BIO Laboratories Inc, CA, USA). No specific permissions were required for sampling in either location nor were any endangered or protected species involved.

### Primer and barcodes design

Primers targeting the V3 and V4 regions of the ribosomal RNA gene originally designed for pyrosequencing (8) were adapted to Illumina sequencing by complementing both Bakt_341F (**CCTACGGGNGGCWGCAG**) and Bakt_805R (**GACTACHVGGGTATCTAATCC**) with sample-specific barcodes. This region of the rRNA gene appears optimal for interrogating bacterial communities (26)

The 7 bp long barcodes were designed to avoid homopolymers and to make two barcodes differ by at least 2 bp. We tested the sequences for self-complementary using Primer3 (http://primer3.sourceforge.net/) and BLASTN. Hairpin loops and self annealing bases were discarded leaving us with 324 identified acceptable pairs of which 50 pairs were randomly selected. In total, 50 barcoded forward primers and 50 barcoded reverse primers where ordered from Eurofins (Germany). A detailed list containing the sequences of the barcodes can be found in supplementary table S1.

### Amplification and barcoding

The variable regions 3 and 4 of the 16S rRNA gene from the single sediment sample was amplified using three different procedures.

i. The starting DNA template material was amplified using non-barcoded PCR primers for 20 cycles (in duplicate and subsequently pooled) followed by a 100 times dilution of the resulting PCR product. Next, in triplicate reproduction, and for every of the fifty barcode pairs, 1 μl of the diluted PCR product was used for 10 additional cycles of amplification with the respective barcoded primers. This batch is referred to as the “two-step PCR” and included 149 samples in three replicate pools.
ii. In singleton reproduction, the DNA material was amplified for 25 cycles directly with the barcoded primers, for each of the fifty barcode pairs. This batch is referred to as the “single-step PCR” and included 49 samples in one pool.
iii. In addition, the starting DNA material from the first pool of the two-step PCR treatment was sequenced a second time in a different run with an updated version of the Illumina chemistry and software. In particular, less random PhiX DNA (5 instead of 50%) was spiked in the sample. This additional dataset is referred to as the “updated chemistry run” and included another 50 samples in one pool. The soda lakes samples were processed accordingly to (iii) and included nine samples which were sequenced in parallel using the 454 technology.

All PCRs were conducted in 20 μl of volume using 1.0 U Phusion high fidelity DNA polymerase (NEB, UK), 0.25 μM primers, 200 μM dNTP mix and 0.4 μg bovine serum albumin. The thermal program consisted of an initial 95 °C denaturation step for 5 min, a cycling program of 95 °C for 40 seconds, 53 °C for 40 seconds, and 72 °C for 60 seconds and a final elongation step at 72 °C for 7 minutes.

To prepare the 248 replicates of the sediment samples for sequencing, the concentration of PCR amplicons was estimated with the Gel Pro analyzer program prior to pooling. Three microliters of each PCR mixture were run on a 1% agarose gel (Invitrogen, Life Technologies Europe BV) in 1X Tris-acetate-EDTA (TAE) buffer stained with gel red dye (0.0001%) and visualized under UV transillumination using a Spectronics variable intensity UV source with diffusor plates and a cooled 12-bit CCD camera (Coolsnap Pro, Media Cybernetics, Silver Springs, MD). The concentration of PCR product was estimated by Gel Pro analyzer 3.1 using 100 bp DNA ladder (Invitrogen) as a molecular weight standard. In the case of the soda lake samples, this quantification was performed ususing a fluorescent stain-based kit (PicoGreen, Invitrogen).

Once combined, every pool of samples resulted in a final amount of approximately 30-40 ng. Following this, the solution was purified by Qiagen gel purification kit (Qiagen, Germany) and quantified using the PicoGreen kit (Invitrogen).

See supplementary figure S1 for an outline of the experimental design.

### Illumina sequencing

The samples were submitted to the SciLifeLab SNP/SEQ sequencing facility hosted by Uppsala University where the routine TruSeq protocol (27) was applied with the exception that the initial fragmentation or size select step was not performed. This involves the binding of the standard sequencing adapters in combination with separate Illumina-specific MID barcodes that enable the combination of different pools on the same sequencing run. This procedure includes an additional PCR totaling 10 more cycles of amplification. As our different barcodes are mixed at this stage, this could potentially cause the formation of chimeric sequences. This protocol also includes the addition of random PhiX DNA to the solution (50%) to provide calibration and help with the cluster generation on the MiSeq’s flow cell. As detailed above, an updated chemistry and less PhiX (5%) was used for some of the samples.

### 454 sequencing

Using the primer pair 341F-805R, the sediment sample as well as nine soda lake samples were processed as described in (28). Sequencing was performed at the SciLifeLab SNP/SEQ sequencing facility at Uppsala University using standard Titanium chemistry.

### Data processing

The data was preprocessed with version 2.1.13 of the Illumina instrument control software for the two-step PCR pools and for the single-step PCR pool. The updated chemistry samples were preprocessed with version 2.2.0. This step includes the separation of pools according to their MID sequence tag and the computation of a few statistics such as GC distribution and quality score distribution. Further statistics were produced using FastQC (29).

#### Barcodes demultiplexing

Using our custom made pipeline, every read-pair produced was parsed and checked for recognizable barcodes on both the forward and reverse sequences. Depending on the outcome of this procedure, read-pairs were classified into one of five categories: 1) The reads which have two different recognizable barcodes at the start of both of the sequences in the pair and these barcodes congruently match to the same sample. 2) The reads which have no identifiable barcodes on either of the sequences in the pair. 3) The reads which have a known barcode on only one of the two sequences of the pair. 4) The reads which have two recognizable barcodes but both barcodes belong to the same family (i.e. we find two froward barcodes or two reverse barcodes). 5) The reads which have two recognizable barcodes but are incongruent and belong to different samples. All categories underwent further processing steps but only the matching barcodes were used in the final results.

#### Assembly

The Illumina technology used does not produce a single character string for every DNA polony on the MiSeq flow cell, but instead produces two strings of fixed size each starting at one end of the original fragment. Fortunately, the length of the 16S rRNA gene region targeted by our primers is short enough to ensure that both sequence ends have to overlap. To recompose the complete nucleotide sequence we used the PANDASeq algorithm (30) at version 2.4. At this step, the overlapping regions were aligned and scored. Alignments that obtained low scores (< 0.6) such as those with short alignment length or high proportion of mismatches were discarded, providing a first step of quality control.

#### Quality control

To further check for erroneous reads, we searched for the forward primer and reverse primer at the start and end of each read, respectively, and discarded those that did not contain them. Following this, we discarded any sequences containing underdetermined base pairs (represented by the letter ‘N’). Furthermore, we scanned every sequence with a sliding window of 10 base pairs and discarded all those that fell below a PHRED score of 5. Finally, we applied a length cutoff and discarded any sequence having an overlap region greater than 100 base pairs.

#### Chimeras

To check for chimeric sequences amongst the different categories of sequences, the UCHIME algorithm (31) included in the free version 6.0.307 of USEARCH was used. Two variations of the program were run and compared. First the denovo mode in which the varying abundances of sequences in the input were exploited. Secondly, we used the reference mode in which decisions are made using a database of chimera-free sequences. For computational time issues, the denovo algorithm was run on 50’000 randomly sampled sequences, while the reference algorithm was run on 100’000 sequences.

#### Clustering

An exact clustering algorithm that computes the difference between every pair of sequences scales with the square of the number of input sequences and hence cannot be used on a dataet of this size. Instead, we used the CD-HIT-OTU package (32) and its variant tailored for Illumina reads (version 4.5.5-2011-03-31). Another heuristic, the UCLUST greedy algorithm (33) (included in the free version 6.0.307 of USEARCH) as implemented in the QIIME (34) script “pick_otus.py” was also tested with default parameters. Thirdly, the latest product from Robert C. Edgar titled UPARSE (16) was applied to our data (included in the free version 7.0.1001 of USEARCH).

#### Taxonomic assignment

For every OTU, the representative sequence of the cluster was used as a query against the quality checked SILVAMOD database using the CREST software (35). This algorithm uses MEGABLAST to quickly search through a hierarchical database of 16S rRNA gene sequences and makes use of a lowest common ancestor strategy to assign each sequence to a particular level of taxonomy in the tree of life. The SILVAMOD database is based on a manual curation of the taxonomical information found in the version 106 of the SILVA SSURef non-redundant release (36). An exception to this procedure was the creation of figure 3 where sequences are not clustered into OTUs and thus resulted in much larger quantities of data. In this case, the RDP naive bayesian classifier version 2.2 was used in combination with its own associated taxonomical database (37).

#### Statistics

Statistical analyses were performed using version 2.0-7 of the VEGAN package (38) and the R statistical framework version 2.11. In particular, these included NMDS ordination plots (**metaMDS()**), beta-dispersion (**betadis()**), PERMANOVA (**adonis()**), permutational ANOVA (**aovp()**) and the estimation of diversity indices. The Bray-Curtis distances are calculated with the usual square transformation and Wisconsin standardization using rarefied datasets.

#### Comparison with 454

The data originating from Roche’s pyrosequencing machines included the sediment as well as 9 soda lakes. In order to maintain comparability, the 454 reads were processed identically to the Illumina data (albeit without the assembly step) through all the steps of demultiplexing, primer presence check, exclusion of undetermined bases, quality filtering and removal of primer sequences. Following these operations the data was pooled with that of the sediment sample and soda lakes sequenced on the Illumina machine. Once combined, all reads were trimmed to 400 bp, clustered using UPARSE and assigned using CREST as described above.

#### Phylogenetic distance

The UniFrac (39) distance was calculated by aligning the representative sequence of every OTU against a 97% clustered version of the Silva SSURef non-redundant database. This database is distributed by the QIIME-group and is based on release 111. The alignment was performed by mothur’s (40) v.1.30.2 align() function with the kmer search strategy and Needleman-Wunsch scoring method. Following this, from the multiple sequence alignment produced, a phylogenetic tree is built with the default settings of FastTree (41) v2.1.7. Finally, the tree and the OTU table are fed into the weighted UniFrac implementation of PyCogent (42) v1.5.3 which computes the final distance matrix.

See supplementary figure S2 for an outline of the data processing. The code produced for the development of this pipeline was written in python and is available at http://github.com/limno/illumitag/ under an MIT license.

The raw sequencing data for the sediment sample and the soda lakes are available at accession numbers SRP044363 and SRP044627, respectively.

## Results

### General performance

As a dummy sample that would be sufficiently complex for our tests, we used material from the upper sediment layer of a Swedish lake. We then ran the sample in multiple replicates and tried to measure the reproducibility of a sequencing experiment on the Illumina MiSeq platform. The total number of paired sequences produced from the dummy sediment sample reached 10‘338’568. Each pair contained two sequences of 250 nucleotides each. The overall PHRED quality scores average was of 36 for the forward reads and of 33 for the reverse reads. As is expected with this technology, the reverse reads were always tagged with a lower quality than the forward reads especially in their terminal region (see figure S3 for more detailed distributions of the quality scoring).

Applying the demultiplexing procedure to all reads revealed that a large proportion of the sequences pairs did not posses matching forward and reverse barcodes. Some sequences did not bear any recognizable barcode at all (6%). Others had only one barcode (14%) placed on either end. Yet another group had two identifiable barcodes but belonging to different samples (22%), thus were not expected to be found together (Figure 1). Within the the remaining 58% that had matching barcodes, proportions of reads were unevenly distributed amongst the barcodes, with a relative standard deviation up to 55%. Figure S4, S11 and S12 detail the distributions of barcodes within each pool. Naturally, only the matching barcode sequences were used in our final analysis and the rest were discarded.

**Figure 1.**
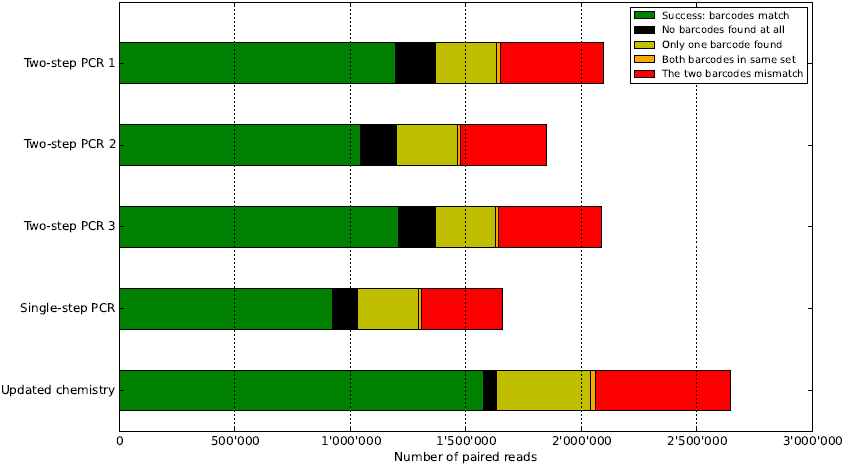
Distribution of barcodes matching and mismatching. To assign an Illumina read to a particular sample, one examines both of the barcodes at each end of the sequence. In green, the two barcodes agree on which sample the read is coming from. In black, no barcodes are found on either end. In yellow, only one barcode is present. In orange, the two barcodes come from the same directional set and should not be found together (e.g. two forward barcodes). In red, the two barcodes each indicate a different sample. Overall, with this setup, about half of the raw data is discarded.

Further sequencing runs not presented in this study were performed using different laboratory quantification methods. Instead of quantifying with Gel Pro, the PicoGreen method was used for every barcode individually. This did not significantly reduce the unevenness of the numbers of sequences per sample (relative standard deviation with pico green 22%, 26%, 88% against 21%, 39%, 49%, 55%, 21% for gel pro).

The assembly procedure, representing the next step in the analytical chain, discarded less than 2% of the matching barcode sequences. Considering the mismatching barcode group, we found that an equal ratio of mate pairs could not be assembled. However, for reads that had none or only one barcode, the unassembled proportions were over 19%. Figure S6 details the distributions of assembled sequences within each group of each pool.

Taking only the matching barcodes group, the length of the overlapping region varied between 20 and 70 bp for the majority of the sequences. A few sequences (0.5%) assembled with an overlap greater than 100 bp and were discarded when applying the length cutoff. Figure 2 shows the distribution of sequence lengths produced which agreed well with the natural variation in the length of the 16S rRNA gene. Three main size fractions appeared, each containing a characteristic species composition. For instance, the fraction between 430 and 446 bp was composted exclusively of Archaea (Figure 3).

**Figure 2.**
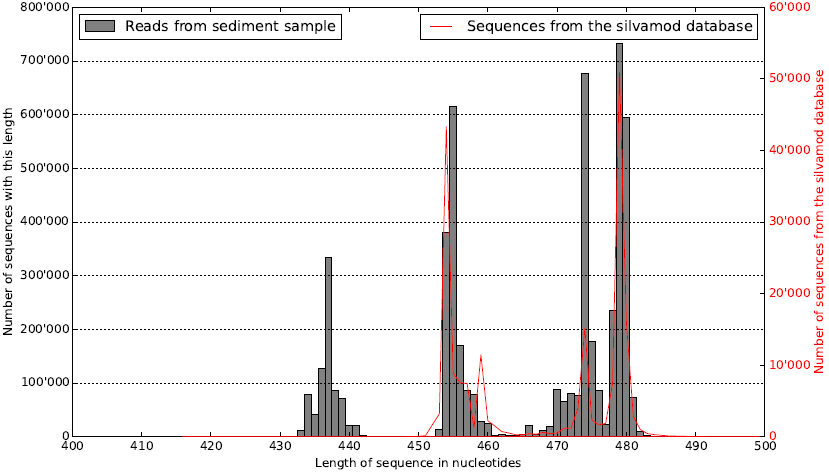
Distribution of sequence lengths for matching barcodes. Once all the forward and reverse reads from the Illumina sequencer are joined one can see – in gray here – a pattern in the size of the fragments produced. Superposing in red the abundance of length variation found in the SILVAMOD database, one can see that the variation we uncovered follows closely the natural variation of the V3-V4 region of the 16S rRNA gene. As shown in figure 3, each of the three
peaks are composed of characteristic phyla.

**Figure 3.**
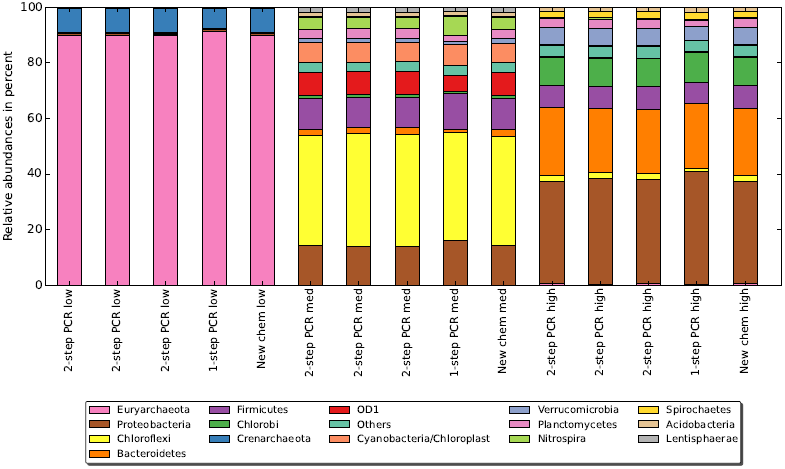
Composition of different lengths fractions. Every one of our sediment sample replicates, i.e. the triplicate two-step PCR, the one-step PCR and the updated chemistry are separated into three size fractions (low, medium, high) according to the peaks identified in figure 2. Fragments below 446 bp are exclusively originating from archaea. The second peak between 447 and 464 bp contains, for instance, contains a majority of the Chloroflexi. The last group above 465 bp holds most of the Bacteroidetes. Other species such as Proteobacteria or Firmicutes are found spanning a size range in the environmental sample.

After assembly, missing primers accounted for an average loss of 4.3% of the remaining sequences. Next, the undermined base pair filtering eliminated on average 0.2%. Following this, the quality control discarded a further 6.0%. This brings the amount of quality filtered reads from our sediment sample to 5’212’432.
Overall, for the reads with matching barcodes, 2.5% were identified as chimeric when using UCHIME in reference mode while up to 23% were identified as chimeric using UCHIME in denovo mode. Taking the fraction of reads with mismatching barcodes these numbers became 5.2% and 36.35%, respectively.

### OTU Clustering

Using the 248 replicates of the sediment sample and applying the CD-HIT-OTU algorithm set with a cutoff at 97% sequence identity resulted in 12’942 OTUs after excluding chimeric sequences. Running the UPARSE algorithm with a minimum difference of 3% between cluster centers produced 14’107 OTUs. In sharp contrast, using the UCLUST algorithm run via QIIME on the same reads and using a sequence similarity threshold at 97% resulted in 189’391 OTUs which is 13 times more than using the default settings in UPARSE.

Once the centroid sequence of each UPARSE OTU was annotated against the SILVAMOD database, all sequences pertaining to the phyla of plastids, mitochondria, thaumarchaeota, crenarchaeota and euryarchaeota were removed. This revealed that 2’152 OTUs representing 16% of the reads were identified as being of non-bacterial origin, which was expected considering the characteristics of the primer-pair.

Keeping the bacterial OTUs and rarefying the number of reads to that of the lowest sample, we can compare the three clustering methods again. After this rarefaction, the UCLUST method resulted in about twice the number of OTUs when compared to UPARSE and CD-HIT-OTU. Indeed, processing the 248 replicated sediment sample in such a manner resulted in, on average, 1’235 OTUs when using UCLUST and, on average, 605 and 830 OTUs when applying CD-HIT-OTU and UPARSE, respectively. The pattern where UCLUST showed much higher numbers of OTUs than the other two methods was also observed when using a separate group of 70 soda lake samples. Yet, plotting OTU accumulation curves (see figure S9) revealed that CD-HIT-OTU and UPARSE followed expected asymptotic trends while UCLUST behaved atypically showing a rise in the amount of rare OTUs. Examining the landscape of assignments on the phyla level revealed that patterns in phylum composition were strongly conserved.

Moreover, beta diversity measures based on Bray-Curtis distances (with sequence numbers rarefaction) applied to a collection of 70 soda lake samples showed highly corresponding trends when comparing the three clustering procedures. Additionaly, a pair-wise Procrustes test among the three OTU tables resulted in coefficients greater than 0.98 and p-values of less than 0.001. Also, trends in evenness were similar and linear models between all three clustering methods resulted in R2 values greater than 0.96 and p-values of less than 0.001. Slopes of these linear models ranged from 0.92 to 1.08 revealing that UCLUST resulted in the most even OTU table followed by UPARSE with CD-HIT-OTU providing the most uneven OTU table. Differences in richness estimated from the three OTU tables corresponded well (R2 ranging from 0.74 to 0.92 and p-values of less than 0.001). The slopes revealed that richness estimates were approximately 40 percent higher with UCLUST as compared to the other two clustering methods. The estimated richness was very similar between CD-HIT-OTU and UPARSE with a slope of 1.03 and an intercept of 27.

### Precision

Choosing the UPARSE clustering method for its speed and simplicity of use, we proceeded to evaluate the precision of the method. The reproducibility of the results and the effects on alpha and beta diversity were determined by comparing results from 248 technical replicates of a single environmental sample run with 50 barcodes in 5 different pools. Permutational MANOVAs on Bray-Curtis dissimilarities revealed significant differences in beta diversity between replicates run in different pools and with different methods. In particular, beta diversity differed significantly between the 1-step and 2-step PCR methods (*R*^2^ = 0.028, p-value < 0.001).

Similarly, using a phylogenetic measure such as UniFrac distances, a significant pool and method effect was observed. The variances in alpha diversity estimates amongst the 248 replicates and the results from the permutational analysis of variance are given in table 1. Here, no significant difference in Chao1 estimates among replicates run in different pools or with different methods or barcodes was observed. However, evenness estimates revealed significant pool and method effects. Applying post-hoc tests revealed that in these latter cases the “single-step PCR” procedure was different from the other four pools. When considering only the two-step PCR pools, no significant difference was found.

**Table 1.**
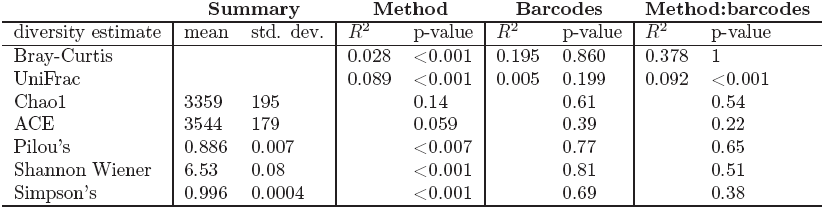
Precision of the Illumina method. The precision of the method over all pools and barcodes is evaluated. For beta-diversity measures, a permutational MANOVA test was used. For alpha-diversity measures, a permutational ANOVA test is used. The effect of methods, barcodes and the combination of methods and barcodes are given.

### Comparisons with 454

Next, we compared 454 pyrosequencing and Illumina sequencing strategies. We evaluated the agreement between the two methods by using 10 samples for which both 454 and Illumina data was available. Reads from the two sequencing machines underwent method-specific quality filtering before being pooled and trimmed to a length of 400 bp. After performing OTU clustering using UPARSE, the consistency in alpha and beta diversity as well as the taxonomic composition was determined. Using Procrustes and Mantel tests, a significant correspondence between beta-diversity estimates was revealed when using Bray-Curtis distances (R=0.995, p<0.001 and R=0.954, p<0.001, respectively). The concordance in beta diversity is also well represented in the dendogram (figure 4) and the NMDS plot (figure S8).

Accordingly, patterns in phylogenetic composition as determined by UniFrac distances also agreed between the two approaches, as shown by Procrustes and Mantel tests (R=0.993, p<0.001 and R=0.968, p<0.001, respectively). We also observed matching results for Chao1 and ACE richness estimates, whereas correspondence was rather low for Pilou’s evenness, Shannon Wiener and Simpson’s index between the two sequencing approaches (table 2).

**Figure 4.**
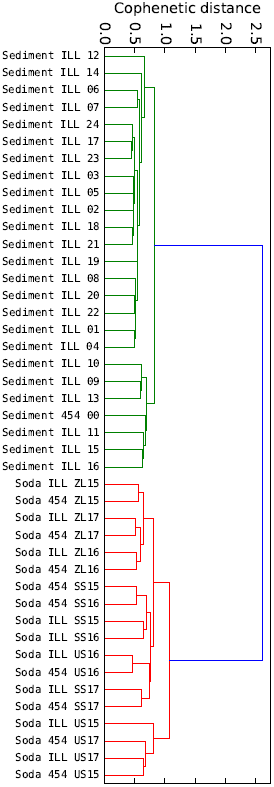
Correspondence of phylogenetic distance between 454 and Illumina. This dendogram included 24 replicates of our test sediment sample sequenced on the Illumina platform as well as the same sample sequenced on a 454 machine. In addition, 9 soda lakes samples that were equally sequenced on both technologies are included. Using the UniFrac metric to compute a distance matrix, a hierarchical clustering is performed following the UPGMA method. The detail of the taxonomic compostion can be seen in figure S7

**Table 2.** Correspondence between 454 and Illumina. A linear model was fitted to the alpha diversity estimates from 454 and Illumina data for each sample. Regressions are plotted in figure S10.

| diversity estimate | $R^2$ | p-value |
| --- | --- | --- |
| Chao1 | 0.99 | <0.001 |
| ACE | 0.99 | <0.001 |
| Pielou's | 0.39 | 0.079 |
| Shannon Wiener | 0.64 | <0.02 |
| Simpson's | 0.17 | 0.196 |

On a taxonomic level, there was substantial overlap in the detected phyla (figure S7). However, the relative phyla contribution was not well conserved between the two methods. The highest discrepancies were observed in samples with substantial proportions of Cyanobacteria.

Performing a paired Wilcoxon test to identify inconsistent OTU abundances between the methods, revealed 18 OTUs with a significance difference (p<0.05). However, false discovery rates indicate that these discoveries are most likely due to chance.

## Discussion

There are several approaches when it comes to amplicon sequencing of the 16S rRNA gene based on the Illumina technology (17) (18) (19) (20) (21) (22) (23). These have been applied to investigate the microbial diversity in numerous environments to great success, even revealing the dynamics of rare taxa.

Here we introduce our own protocol starting with PCR amplification of bacterial 16S rRNA genes, followed by paired-end Illumina sequencing and ending with bioinformatic analyses. Our experimental Illumina tag sequencing design used barcoded primers flanking the V3-V4 segment of 16S rRNA gene, a region commonly amplified in pyrotag experiments (8) (43) (28) (44). To construct a standard Illumina paired-end library with an individual MID, 50 individual samples were amplified, mixed and then used as templates each time. Usually 4-6 of these MID coded libraries were then simultaneously sequenced on an Illumina MiSeq. With the current read length of twice 250 bp, the V3-V4 region of the rRNA gene presents an optimal target for sequencing (26) as it provides an adequate overlap of the forward and reverse paired-end reads. Moreover, assembling these reads increases the quality and confidence in the overlapping region (20) (22).

### Where did all the sequences go?

Applying our bioinformatic pipeline, over half of the paired-end reads from each Illumina run were subsequently discarded. This was due to either (i) low quality score (ii) unassembled pairs (iii) assembled pairs that contained mismatched barcodes (iv) sequencing errors in one or both of the primer regions
(v) archaeal or eukaryotic sequences. Other Illumina-based 16S rRNA gene studies have encountered similarly high error rates, resulting in such extreme read filtering (approximately by a factor 2).

Similar to our results, high incidence of mismatching barcodes (19) have been previously reported as the main loss factor. In this earlier study, over-clustering and sequence chimeras were ruled out, and instead primer contamination during the initial sample amplifications were given as the most likely cause. Laboratory contamination is always a possibility that can explain the mismatches. However, in our case, this is unlikely as the high proportion of barcode mismatches remained unchanged in every experiment conducted for this study as well as in all the following sequencing runs that are not presented here. Indeed, multiple individuals have reproduced the protocol presented in this article and all obtained similar results with regards to sequence loss.

A technical issue within the Illumina machine could be another source, such as erroneous identification of the DNA clusters on the flow cell by the imaging software. Comparing matching and mismatching barcodes showed that assembly performed constantly in both cases permitting us to refute the hypothesis that the mismatching barcodes are due to the paired end sequencing. Furthermore, reports of such problems are not prevalent and the metrics generated show that at least 90% of the clusters passed the flow cell filtering algorithm for the first four pools and at least 85% for the last pool.

Our present interpretation and argument is that mismatched barcode sequences are most likely produced in the library preparation. Our experimental design obviously amplifies each sample with its corresponding barcodes separately. However, there is a supplementary PCR performed by the sequencing facility, occurring after all samples are pooled together and adaptors are added. We hyothesis that this causes the chimeras which is supported by the chimera detection results. Indeed, both algorithms identified that the mismatching barcodes group had a slightly larger proportion of chimeras than the group with matching barcodes. Yet, the increased proportion of chimeras in the mismatching barcodes group is rather low and is not sufficient to entirely explain the proportion of mismatches. Most likely, chimeric sequences are formed during the Illumina library preparation by highly similar amplicons that originate from differently barcoded samples. This kind of “perfect” chimeras could be the ones that go unnoticed. It is also worth mentioning that, as a proofreading polymerase is used in the library preparation, there is a risk of unstable amplification. Such polymerases can fall off from time to time creating partial amplicons which will be used in “false” priming to produce “perfect” chimeras. Still, the proportions of chimeric sequences varied greatly depending on the algorithm used and cast doubts on the specificity of the chimera detection algorithm.

A possible solution might be provided by using a primer construct including Illumina adapters and 16S rRNA gene specific primers in the first step PCR, combined with the attachment of standard Illumina handles and index primers in the second step PCR. This represents the next generation procedure, already under development in our lab, where PCR amplification and library preparation will be combined.

A secondary issue is the unevenness of the read coverage produced per sample. If every barcode represented a unique environmental sample, unlike in our current evaluation experiment, one would typically prefer the quantity of data produced for each sample to be equal. This is for example the case if one wants to compute statistical measurements that are sensitive to sample size, where one is forced to rarefy the read counts to the lowest sequence group. Sources for the unevenness are pipetting errors and uncertainties in quantification for which we have no solution as different procedures all performed equally “unsatisfactory".

Another interesting observation following our study is that the length distribution agrees with the natural variation in the lengths of the 16S rRNA gene. Such length polymorphisms effects quantification as shorter reads are known to be preferentially amplified and sequenced, which also suggests that observations of multiple bands on electrophoresis gels of bacterial community 16S rRNA gene amplicons are not an artifact of PCR.

### OTU making and diversity estimates biases

We observed major differences in absolute richness estimates, and minor differences in evenness which can be explained by the heuristics of the clustering algorithms. Thus, alpha diversity estimates based on OTU tables obtained by different clustering methods should not be compared without correcting for general discrepancies. Such corrections can be performed using linear models as obtained in our study. The applicability of such linear models is supported by our set of 9 soda lake samples, but the universality of the models requires further evaluation.

Besides the differences in absolute estimates, the high R^2^ and p values of the linear regression analyses reveal that general trends in alpha and beta diversity were conserved and corresponded well regardless of the clustering algorithm. While all clustering methods revealed conserved and corresponding trends, we recommend the usage of UPARSE because this algorithm requires the lowest CPU allocation.

Taking a closer look at the alpha diversity in the 248 replicates, relative standard deviation never exceeded 5% among replicates, while average estimates of richness determined from all replicates were approximately 3.6 times lower than the numbers of OTUs detected in all replicates. This underestimation of richness as well as the variations among replicates are most likely due to sampling artifacts associated with random sampling (45), as well as the performance of the technology per se. Many steps in the Illumina tag sequencing procedure are associated with random sampling, including PCR amplification of target genes, ligation of amplified PCR products to sequencing adaptors, amplification of ligation products and immobilization to flow cells, as well as identification of the DNA clusters on the flow cell by the imaging software. One way to improve reproducibility and quantitation is to use biological replicates also when considering upstream procedures such as environmental sampling and DNA extraction.

### Can 454 and Illumina data be combined?

Picking 24 Illumina sequenced replicates of our sediment sample and its corresponding 454 run as well as the nine soda lakes samples which were both run with Illumina and 454, we obtained highly corresponding trends in richness and beta diversity but not in evenness. A possible explanation for the lower evenness produced by the Illumina technology could be the presence of the additional PCR step in the library preparation.

Computing the UniFrac metric on all the samples, we note that the average distance amongst samples increases as one moves from a series of replicates using the same technology (mean distance amongst Illumina sediment replicates 0.404) to a replication of the same sample with two different technologies (mean distance between 1×454 sediment and 24×Illumina sediment 0.499) to a group of samples taken from similar environments (mean distance between all soda lakes 0.686) and finally to the comparison of two totally different environments (mean distance between Illumina sediment and Illumina soda lakes 0.812). These trends hold up when using the Bray-Curtis distance metric.

Comparing 454 and Illumina by using the same PCR primers and bioinformatic analyses resulted in corresponding trends in richness and beta diversity. Nonetheless, taxonomic composition was proportional biased which is also reflected in the non corresponding patterns in evenness.

## Conclusion

The main conclusion of this report are: (i) Mismatching barcodes between forward and reverse reads, indicative for chimeras, are most likely introduced in the amplification step of the library preparation. Thus, reverse and forward primers need to be complemented with unique barcodes to avoid miss-assignment of reads to samples when employing the standard TruSeq library preparation protocol. (ii) Although, different clustering algorithms result in different numbers of OTUs, trends in alpha and beta diversity are conserved. (iii) For those switching sequencing technologies, 454 and Illumina sequence data can be combined provided the following conditions are respected: the same PCR primers and bioinformatic workflows must be applied and the variations between the methods must be quantified and accounted for in the interpretation of the results.

## Acknowledgements

This work was supported by the Swedish Foundation for Strategic Research (Grant Number ICA10-0015 to AE). The work was also funded by a postdoctoral stipend from the Carl Tryggers Foundation (to OA) and a grant from the Swedish Research Council (to SB). We also would like to acknowledge the support for the 454 sequencing made possible via an instrument grant from the K&A Wallenberg foundation and the support for the Illumina sequencing made by the SciLifeLab SNP/SEQ facility hosted by Uppsala University. The computations and bioinformatics were performed on resources provided by the Swedish National Infrastructure for Computing (SNIC) through Uppsala Multidisciplinary Center for Advanced Computational Science (UPPMAX) under Project “b2011032". Finally, we want to thank Alexander KT Kirschner and Stefan Jakwerth for their assistance with sampling of the soda lakes in Austria, as well as Sainur Samad for his assistance with DNA extraction and processing in the laboratory.

